# Embracing firefly flash pattern variability with data-driven species classification

**DOI:** 10.1101/2023.03.08.531653

**Authors:** Owen Martin, Chantal Nguyen, Raphael Sarfati, Murad Chowdhury, Michael L. Iuzzolino, Dieu My T. Nguyen, Ryan M. Layer, Orit Peleg

## Abstract

Many nocturnally active fireflies use precisely timed bioluminescent patterns to identify mates, making them especially vulnerable to light pollution. As urbanization continues to brighten the night sky, firefly populations are under constant stress, and close to half of the species are now threatened. Ensuring the survival of firefly biodiversity depends on a large-scale conservation effort to monitor and protect thousands of populations. While species can be identified by their flash patterns, current methods require expert measurement and manual classification and are infeasible given the number and geographic distribution of fireflies. Here we present the application of a recurrent neural network (RNN) for accurate automated firefly flash pattern classification. Using recordings from commodity cameras, we can extract flash trajectories of individuals within a swarm and classify their species with a precision and recall of approximately seventy percent. In addition to scaling population monitoring, automated classification provides the means to study firefly behavior at the population level. We employ the classifier to measure and characterize the variability within and between swarms, unlocking a new dimension of their behavior. Our method is open source, and deployment in community science applications could revolutionize our ability to monitor and understand firefly populations.

## 1 Introduction

Nocturnal fireflies (Coleoptera: Lampyridae) have evolved a visually impressive light-based communication system under simultaneous evolutionary pressures to advertise their species, accentuate their sexual fitness, and avoid predation [1–3]. Using a luciferin-luciferase reaction within their abdomen [4], fireflies broadcast their species identity via light pulses [1, 5]. In many North American genera both sexes flash and must also encode biological sex in their signals [1, 6, 7]. Sympatric species must also produce distinguishable patterns to effectively communicate species identity [1, 5, 8].

This unique signaling system makes fireflies particularly susceptible to human-created population threats like light pollution [9–12]. Artificial light at night (ALAN) interferes with fireflies’ perception of conspecific signals and disrupts their communication timings, curtailing flash signaling behavior and preventing successful mating [10, 13]. In addition to light pollution, habitat degradation, pesticide use, water pollution, and climate change comprise some of the most serious environmental stressors causing declines in firefly populations across the globe [14]. Recent Red List assessments by the International Union for Conservation of Nature (IUCN) have identified that at least 14% of 132 firefly taxa in the United States alone are in danger of extinction [12], but this value is a likely underestimate due to a lack of information on over half the species assessed [6, 12]. Further studies and fieldwork are urgently needed to monitor changes in firefly abundances and evaluate and mitigate the threats imperiling fireflies worldwide [6, 12]. Many other insect species share these environmental threats and are experiencing dramatic declines [10, 15, 16]. Because of their charismatic nature and popular appeal, fireflies can serve as a flagship symbol to foster public attention toward this conservation crisis.

The major barrier to planning effective conservation efforts is the dearth of quantitative data on firefly populations [6, 12, 14, 17]. The high-throughput population monitoring studies that provide the fundamental data for understanding population-level dynamics [18] have not been performed for nearly all firefly species [6]. This gap is largely due to limitations of existing monitoring methodologies, which require the presence of human expert observers [6], are often subjective [6], and typically characterize firefly flash behavior by single, imprecise pictorial representations despite known temperature dependence and individual variability [7, 8]. To address the data deficiency in current conservation efforts, we propose a scalable, automated population monitoring method that combines recent advances in stereoscopic filming, computer vision, and machine learning to classify individuals and quantify swarm-level dynamics. Our method starts with a field recording of a swarm using two consumer-grade cameras (Fig. 1A), produced following procedures described previously in [19]. From this recording, we identify individual firefly trajectories and extract their flash-pattern time series (Fig. 1B). We then trained a recurrent neural network (RNN) on these data to accurately determine species identity from nothing more than the temporal differences in each species’ flash pattern, achieving precision and recall of approximately 0.70 (Table 1). We additionally visualize the distinguishability of firefly flash patterns via a dimensionality reduction of the weights of the last hidden layer of the neural network, using t-distributed stochastic neighbor embedding (t-SNE) [20]. The t-SNE embedding reveals significant clustering by species, but multiple species can coincide in the same cluster, illustrating the intra-species variability that contributes to the difficulty of the problem (Fig. 1C). To our knowledge this is the first application of machine learning to firefly behavioral biology.

**Figure 1.**
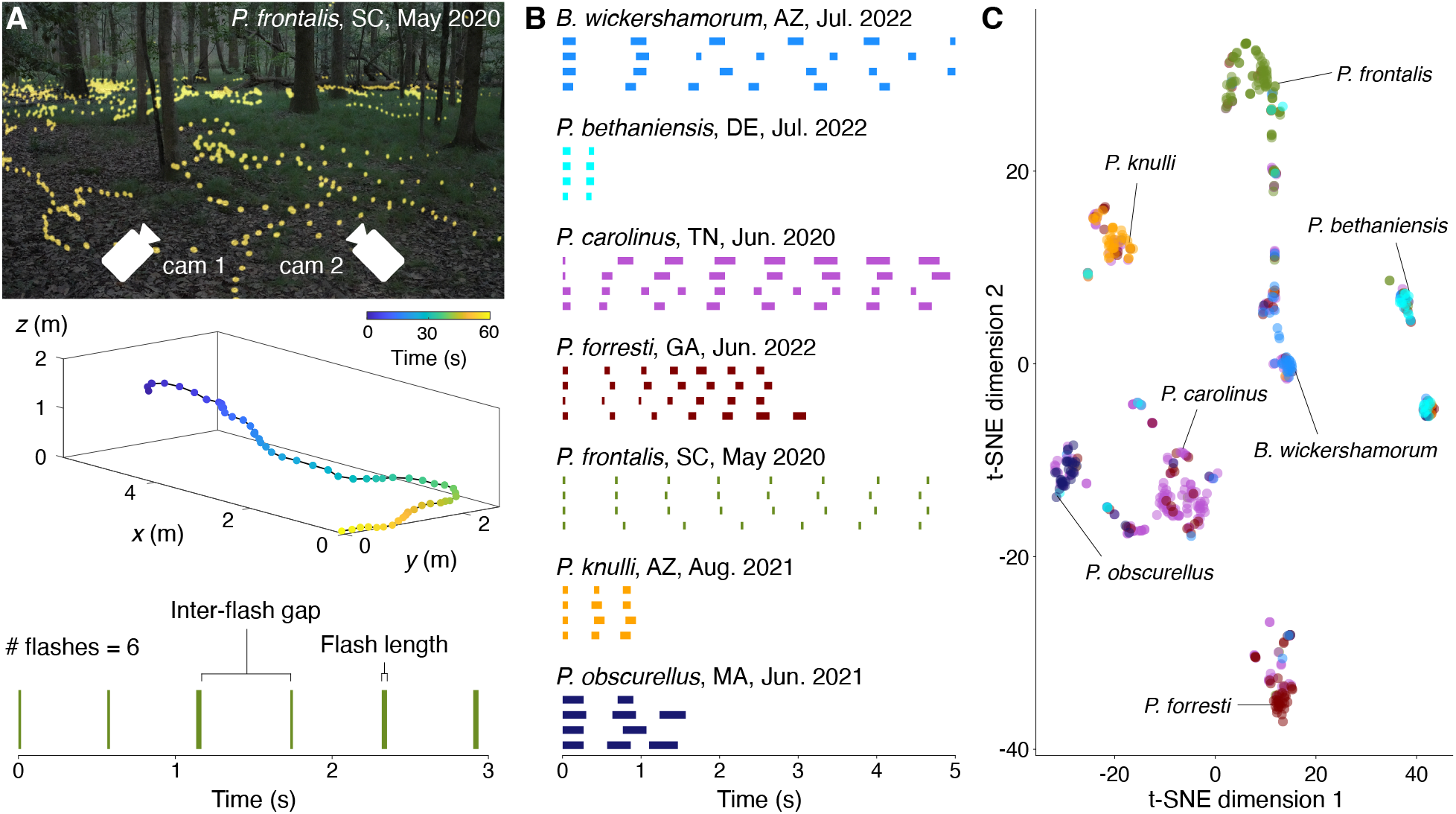
**(A)** Our standardized data collection method **(top)** films fireflies in their natural habitat with two cameras arranged in a stereoscopic vision configuration [22] (photo reproduced and modified with permission from [23]). Flash streaks (**center**, colored by time) in the resulting videos are triangulated and concatenated into trajectories, based on proximity and velocity. (**bottom**) Each trajectory is represented by a time series of individual flashes that can be parameterized with three values: the number of flashes; the flash length, or the average amount of consecutive time for which a flash is detected; and the inter-flash gap, the average amount of time from the end of a flash to the start of another flash. **(B)** Four example five-second flash sequence time series for each of the seven species in our study, labeled with the location and date of the corresponding recording. Our recurrent neural network is used to classify flash sequence time series; time series shown were selected from the top 100 sequences per species with the highest classification probabilities following the filtration process outlined in Methods section 4.4. **(C)** Two-dimensional t-SNE embedding of the output of the last hidden layer of the network, just before inference. Flash patterns are clustered by similarity, and the distinguishability of each species’ characteristic flash pattern can be detected by the colored clusters. However, overlapping points do occur due to the variability of species’ flash signals and the precision of the data-acquisition methodology (see Methods).

Using the most probable predictions of each species, we also provide estimates of flash phrase variability, including the first-ever quantitative characterization of *Bicellonycha wickershamorum* and *Photuris forresti*, two formerly data-deficient species which may be severely threatened [21]. Automated and data-driven methods like the one proposed here are essential to scaling ecology and conservation biology projects [17]. This work enables accurate identification and classification of firefly species in order to ensure their protection and long-term survival.

## 2 Results

Using our precise, high-throughput data acquisition method (Fig. 1A and Methods section 4.1), we recorded nine firefly swarms during the summer months of 2020, 2021, and 2022 that comprise seven North American firefly species and total 38,081 flash pattern time-series after cleaning (see Methods section 4.1 and [24] for the original dataset). These data are a dramatic expansion from previously published results, which were primarily single characteristic patterns per species, and represent firefly swarms filmed each summer over the past three years. This large dataset enables us to leverage machine learning for species classification and characterization of intra-swarm variability. Using an 80:10:10 train:validation:test split of these data, we trained a recurrent neural network (RNN) to produce class label predictions based on the temporal information in each sequence (see Methods section 4.2.1). The accuracy of these predictions was compared to those obtained from several alternate classifiers utilizing signal processing metrics (Table 1 and Methods section 4.3.3). We present ensemble results evaluating the model’s performance on the test set for each method in Table 1. The RNN achieves the best performance in both categories, with an increase of approximately 20% and 60% higher precision and recall, respectively, over the second best classifier. The RNN classifier additionally generalizes across filming sites and filmographers, demonstrating no observable batch effect in its accuracy to the limit of the data we have collected so far (see Supplementary section 1). This importantly enables us to have confidence in the model’s classification decisions even as we continue to train the model in the future, adding new populations of existing species and updating the model’s knowledge without impacting accuracy.

**Table 1:** Summary of signal classification methods and their performance on the firefly flash pattern dataset. ± values indicate standard error of the mean for each statistic.

| Metric | Precision |  | Recall |  |
| --- | --- | --- | --- | --- |
|  | Literature | Population reference | Literature | Population reference |
| SVM | 0.34 | $0.36 \pm 0.00084$ | 0.30 | $0.53 \pm 0.0014$ |
| Jaccard index | 0.23 | $0.41 \pm 0.0011$ | 0.26 | $0.48 \pm 0.0019$ |
| Dot product | 0.24 | $0.25 \pm 0.0025$ | 0.25 | $0.38 \pm 0.0013$ |
| Dynamic time warping | 0.27 | $0.43 \pm 0.00040$ | 0.27 | $0.58 \pm 0.0013$ |
| RNN | **$0.70 \pm 0.0019$** |  | **$0.71 \pm 0.0019$** |  |

We first attempt to utilize published, pictorially represented flash patterns (see Methods section 4.3.1)to classify sequences with signal processing methods. We constructed three different classifiers that use the Jaccard index, dot product, and dynamic time warping (see Methods section 4.3.3) to compare our time series data to the literature references. We additionally constructed a support vector machine (SVM) that classified the time series data from the three flash pattern parameters (flash length, inter-flash gap, and the number of flashes) (Fig. 1A, bottom). These literature-based classifiers perform poorly, with the SVM achieving the highest precision and recall of 0.34 and 0.30, respectively (Table 1).

We attribute the poor performance of the literature reference methods to the ineffectiveness of a single reference sequence per species at capturing the intra-species variability inherent to a large dataset. To address this, we create “population reference” sequences by aggregating sequences of each species in our dataset, aimed at capturing the intra-species variability while preserving species-specific characteristics of the flash signal. We perform an 80:20 train:test split, generate population references from the training set, and classify the remaining test data. The performance of all four classifiers improves slightly by using population references, with dynamic time warping achieving the highest precision and recall of 0.43 and 0.58, respectively (see Table 1). Additionally, we note the literature are incomplete with regard to endangered species such as *B. wickershamorum*, one of the species most represented in our dataset.

In Fig. 2 we show confusion matrices for each ensemble to further illustrate the per-class capabilities and shortcomings of each method. While the dynamic time warping and dot product classification methods occasionally produce a high scoring class (see Methods Fig. 5F), the RNN results reveal a balanced, highly accurate classifier regardless of species, and as such is our recommendation for the algorithm of choice moving forward in this space. Per-species precision and recall metrics can be found in Methods Fig. 5.

**Figure 2.**
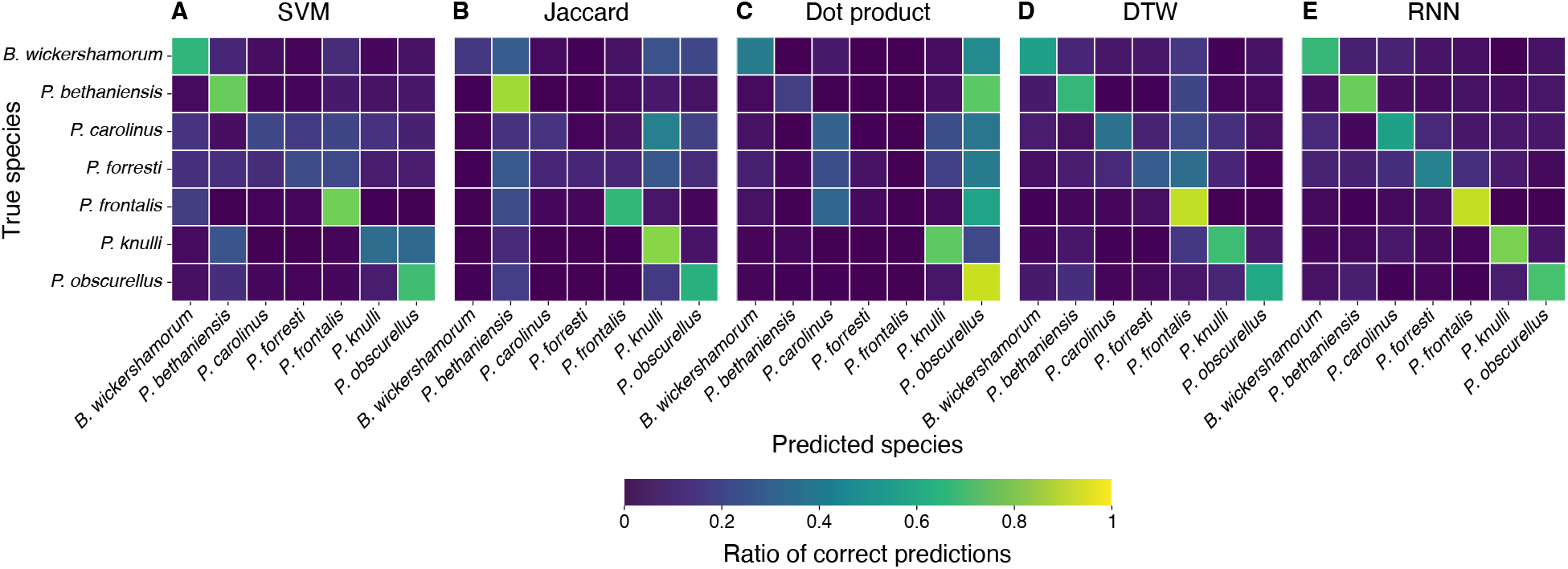
Confusion matrices for each classification method. **(A-E)** Ratios of true positives and false negatives along the horizontal, true positives and false positives along the vertical. Each square in the diagonal represents the recall for the class, and brighter colors indicate higher values as shown in the colorbar. A perfect classifier would consist of yellow squares at each diagonal position and dark purple off the diagonal.

By combining our extensive data set and effective RNN, we provide deeper insights into firefly behavior with a data-driven characterization of a swarm’s signaling pattern. For each swarm recording, we aggregate the one hundred most confidently classified flash patterns (see Methods section 4.4) to produce empirical distributions for flash length, inter-flash gap, and the number of flashes for each species (Fig. 3). These distributions represent the first known quantification of firefly behavioral variability from data. Table 2 compares the mean and standard deviation of flash parameters from our data with the published literature values, where available. Encouragingly, all the previously published numbers of flashes were statistically indistinguishable from our distributions. Half of the published inter-flash gaps and all the published flash lengths differed significantly from our distributions. These differences may be due to improved measurement accuracy (see below) or could represent behavioral differences between swarms, motivated by a changing climate or other environmental factors.

**Table 2:** Mean and standard deviation of the number of flashes, inter-flash gap, and flash length from the filtered data for each filmed population, along with the corresponding literature values (where available) extracted via methodology shown in Fig. 4.

| Species | Number of flashes |  | Inter-flash gap (s) |  | Flash length (s) |  |
| --- | --- | --- | --- | --- | --- | --- |
|  | Data | Literature | Data | Literature | Data | Literature |
| *B. wickershamorum*, AZ, Jul. 2022 | $3.9 \pm 1.1$ | n/a | $0.79 \pm 0.08$ | n/a | $0.14 \pm 0.03$ | n/a |
| *P. bethaniensis*, DE, Jul. 2022 | $2.1 \pm 0.3$ | 2 | $0.13 \pm 0.11$ | 0.53 | $0.06 \pm 0.03$ | 0.27 |
| *P. carolinus*, TN, Jun. 2020 | $7.1 \pm 2.1$ | 9 | $0.44 \pm 0.08$ | 0.37 | $0.07 \pm 0.02$ | 0.23 |
| *P. carolinus*, OH, Jun. 2022 | $5.5 \pm 1.1$ | 9 | $0.43 \pm 0.04$ | 0.37 | $0.09 \pm 0.02$ | 0.23 |
| *P. forrestii*, GA, Jun. 2022 | $6.6 \pm 1.8$ | 5 | $0.39 \pm 0.09$ | 0.10 | $0.09 \pm 0.04$ | 0.16 |
| *P. frontalis*, SC, May 2020 | $13.1 \pm 6.1$ | 10 | $0.56 \pm 0.06$ | 0.87 | $0.04 \pm 0.004$ | 0.13 |
| *P. frontalis*, TN, Jun. 2021 | $10.9 \pm 3.9$ | 10 | $0.59 \pm 0.12$ | 0.87 | $0.04 \pm 0.01$ | 0.13 |
| *P. knulli*, AZ, Aug. 2021 | $2.5 \pm 0.6$ | 3 | $0.35 \pm 0.03$ | 0.32 | $0.07 \pm 0.02$ | 0.20 |
| *P. obscurellus*, MA, Jun. 2021 | $2.8 \pm 1.1$ | 3 | $0.34 \pm 0.10$ | 0.28 | $0.25 \pm 0.11$ | 0.22 |

**Figure 3.**
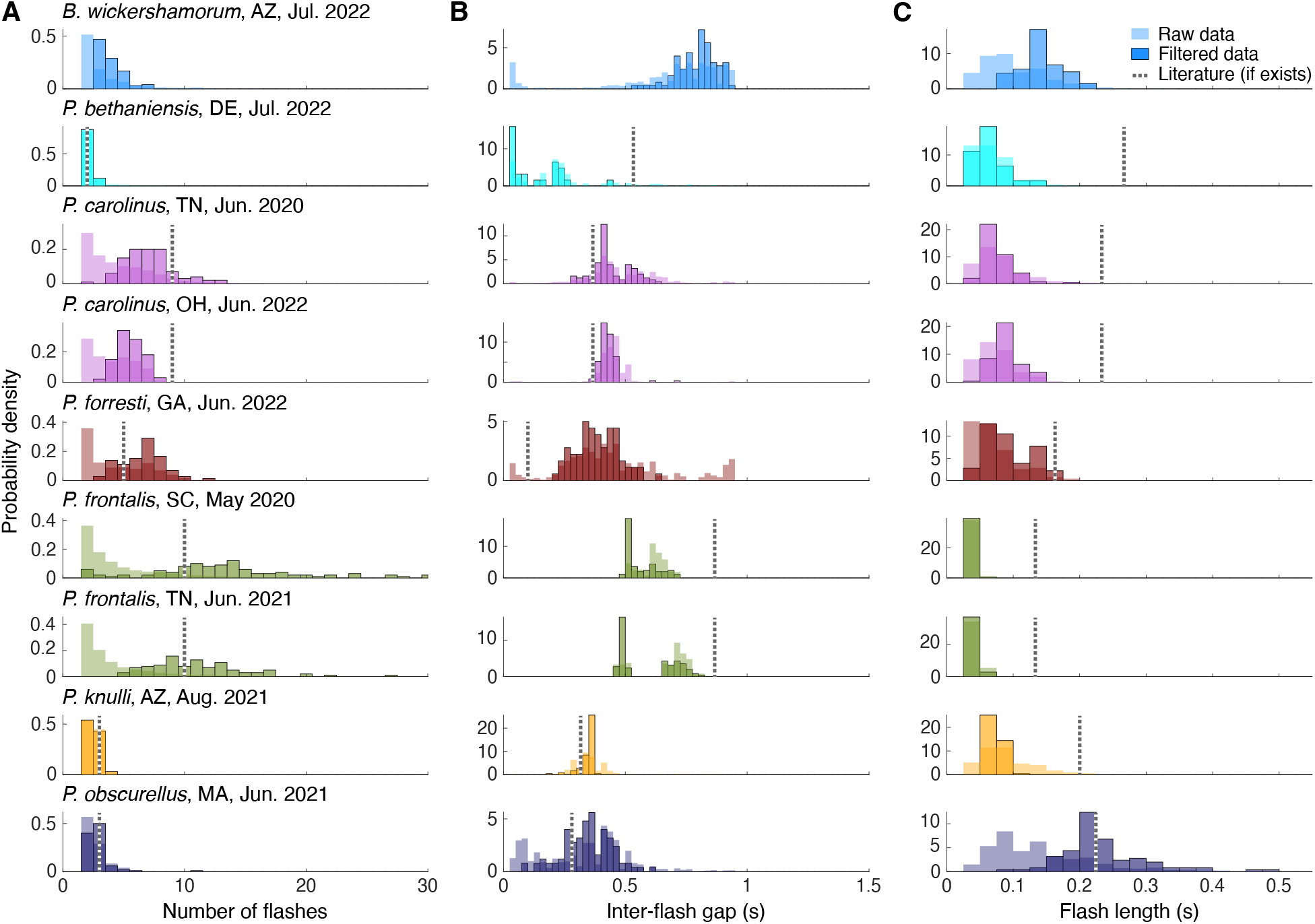
Data-driven characterization of flash patterns for each filmed population, encompassing 7 different species. *P. carolinus* and *P. frontalis* data were each collected from two populations filmed in different locations and years, each of which is separately characterized here. For each population, the 100 sequences with the highest classification probabilities are used to characterize flash signals. This represents a data filtering procedure that extracts the most salient properties of each population’s flash behavior while retaining inherent intraspecies behavioral variability. Probability distributions of the **(A)** number of flashes, **(B)** inter-flash gap in seconds, and **(C)** length of flash in seconds are shown for each species, normalized to sum to 1 under the interval, for both the raw (transparent bars) and filtered (opaque bars) data. The corresponding values obtained from the literature references (see Methods section 4.3.1), if they exist, are shown as dashed lines.

**Figure 4.**
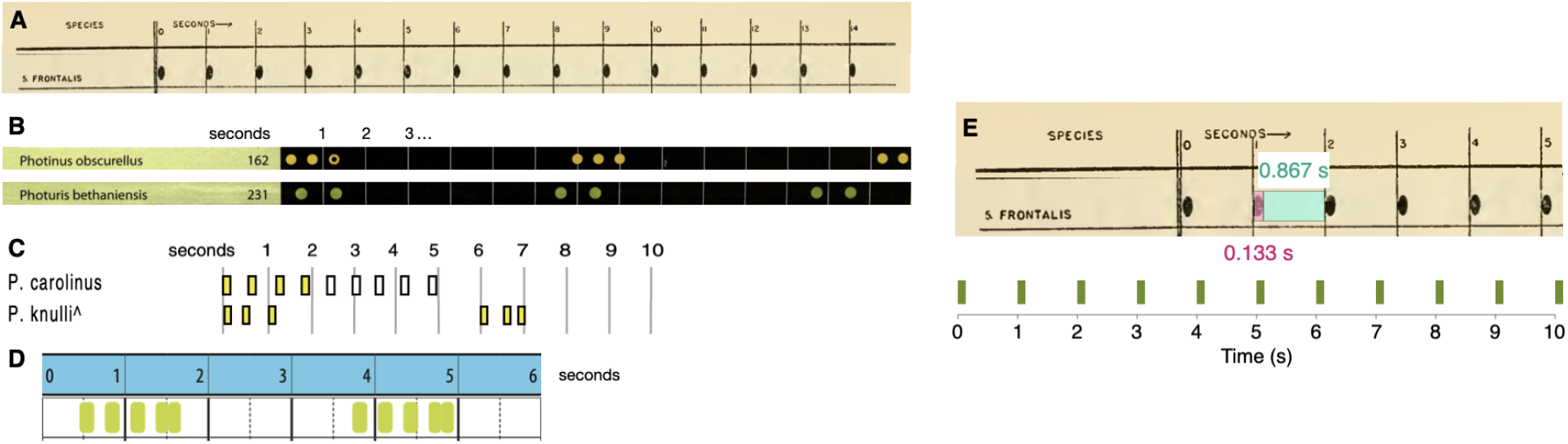
Reference sequences for the firefly species examined in this paper, as previously published in the literature, with the exception of *B. wickershamorum*. **(A)** Reference pattern for *P. frontalis* from Barber, 1951 [42]. **(B)** Reference patterns for *P. obscurellus* and *P. bethaniensis* from Faust, 2017 [7]. **(C)** Reference patterns for *P. carolinus* and *P. knulli* from Stanger-Hall and Lloyd, 2015 [8]. **(D)** Reference pattern for *P. forresti* from Fallon et al., 2022 [6]. **(E)** Illustration of extracting *P. frontalis* sequence from literature pattern (top) and converting to time series (bottom).

**Figure 5.**
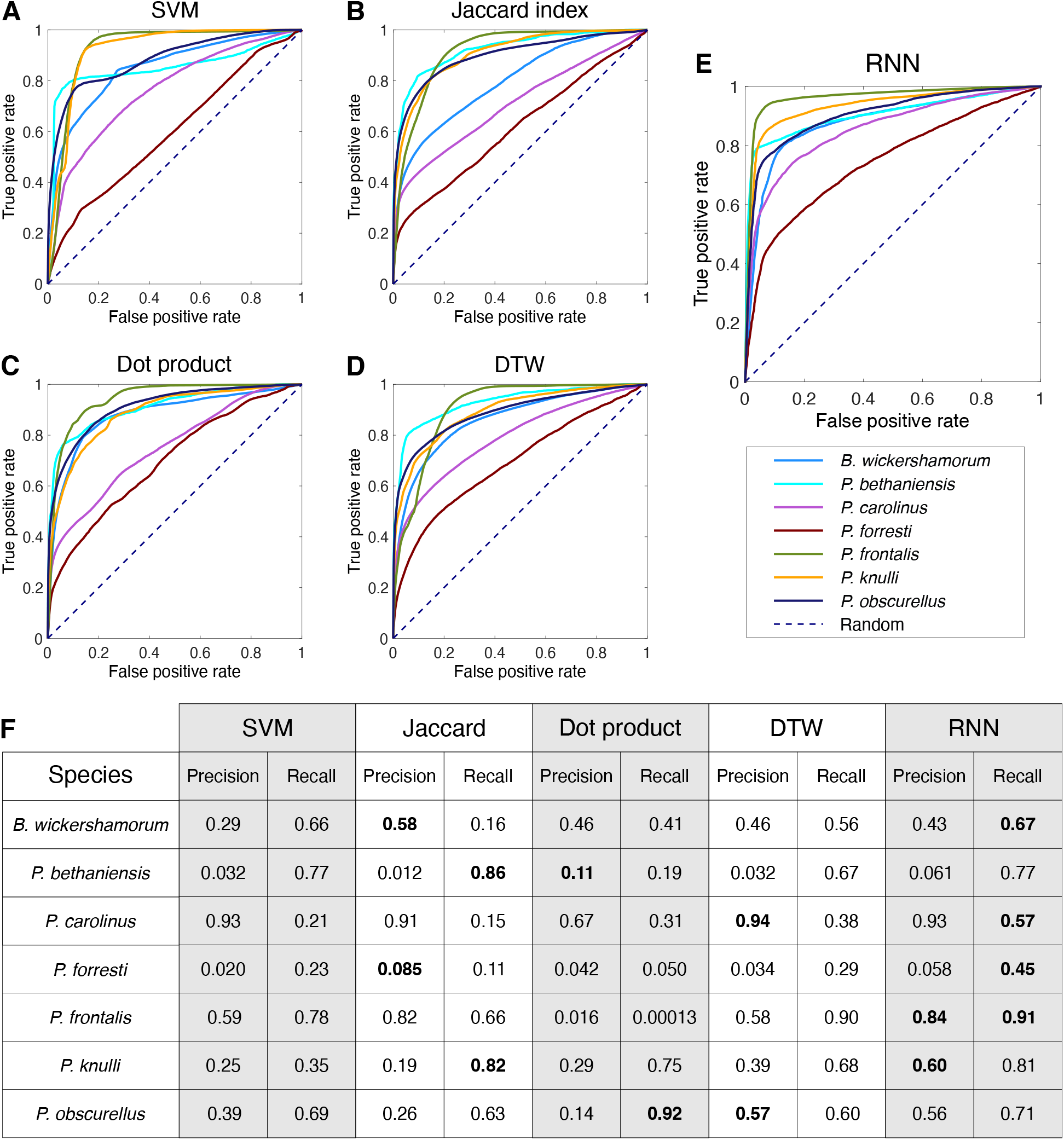
Per-species ensemble of classification results. **A-E**: Receiver operating characteristic (ROC) curves representing the true positive rate (TPR) as a function of the false positive rate (FPR) across all model thresholds of classification, labeled by method. All non-RNN classification methods are conducted using population references. **F**: Table of per-species precision and recall across all surveyed methods (N=100). Bold statistics in the table represent the highest performer for each metric and species. Precision values for *P. bethaniensis* and *P. forresti* are low because these two species represent the classes with the fewest number of samples, and so there is a very small amount of true positive values. However, these still greatly exceed what would be expected by chance (0.001 and 0.006, respectively). The high recall for these classes indicates that the true positives are correctly captured.

These data-driven distributions have several advantages over manual timekeeping. Firstly, representing a species’ flash-based communication as distributions of signal parameters rather than a single pattern allows us to explore the extent of the temporal variability present in a population or across populations. We observe, for example, high variability in the number of flashes emitted by *P. frontalis*, while the flash trains of *P. knulli* exhibit very low variability in the number of flashes (Fig. 3A). Meanwhile, both of these species display relatively tight distributions of inter-flash gap (Fig. 3B), suggesting that these species require precise timing of flashes for communication.

Our methods are more precise than manual timing in measuring the flash length of short flashes perhaps due to the cameras’ high temporal resolution (30 Hz). Our results reveal that the flash length distributions for five species are shifted considerably toward shorter flash lengths than the published literature results (Fig. 3C); this discrepancy may potentially be a consequence of the lower temporal resolution inherent in manual timekeeping due to system lag.

Overall, the filtering process that produces data-driven characterizations of flash patterns in Fig. 3 preserves the most salient characteristics of each species’ signaling behavior while also illustrating the intraspecies behavioral variability. This variability may be substantial in only some of the 3 parameters: for example, *P. obscurellus* demonstrates low variability in the number of flashes but the largest variability in flash length out of the 7 species studied. In comparison, the flash lengths of *P. frontalis* sequences are all extremely similar, but the number of flashes in a train can vary greatly.

We note that these characterizations only represent the locations and conditions where the underlying data were recorded; these probability distributions may be different for data taken in different habitats, at different temperatures, or at different times. By deploying our data acquisition procedure to more sites, we will be able to better characterize the rich extent of a firefly species’ communication.

Previous studies have examined the effect of environmental factors such as artificial light at night (ALAN) on firefly signaling by quantifying the reduction in flash rate or the number of flashing individuals [10]. Our method of generating data-driven characterizations will enable a more acute investigation of how ALAN and other factors affect firefly flash pattern parameters down to an individual level. In addition, we can also investigate whether responses to environmental factors are encoded in behavioral variability: it is unknown whether the variability in flash behavior changes with light pollution and the resulting implications for firefly signal processing and mating success.

## 3 Discussion and future directions

There are 171 known firefly species in North America [6], many of which are endangered and/or lack quantitative data on their flash patterns and intra-species behavioral variability. These knowledge gaps significantly limit our ability to comprehend and respond to environmental pressures. By integrating an inexpensive data-gathering procedure with the presented classification method, we now have the core components of an automated swarm monitoring system that aligns with outcomes suggested in [17]. This can provide crucial data for maintaining the amazing biodiversity within the *Lampyridae* family, preserving the wonder and fascination of firefly bioluminescence for generations to come.

The affordability and portability of the filming system makes it ideal for deployment by researchers and citizen science volunteers [24] to firefly hot spots across the continent. Known species can be monitored for behavioral or population density changes by comparing new observations to historical data, even when considering swarms whose species’ baseline characterization was established at a different location or temperature. For example, despite the variability in inter-flash gaps, our model produced accurate classifications for *P. frontalis* swarms collected at different sites, by different individuals, and under different conditions (Supplementary section 1). New species can be added to the growing corpus once sufficient trajectories are acquired such that the neural network can be retrained. From this work, we estimate that at least one hundred trajectories per species are required before the model can robustly distinguish their patterns, and these can be acquired from just a few days of filming. Going forward, all species should be targeted for future deployments to establish their flash dynamics baseline and continue with periodic monitoring. However, the most critical and immediate need is to deploy cameras to areas containing critically endangered or vulnerable firefly species, such as *B. wickershamorum, P. bethaniensis*, and *P. knulli*.

Our system embraces intraspecies variability by producing data-driven characterizations of firefly flash patterns with local granularity. These characterizations align well with known descriptions, where data are available, and provide statistical clarity for several endangered species for which data are scarce or non-existent: our flash sequence characterization of *B. wickershamorum* and *P. forresti* represent the first known quantitative description of the flash pattern for these vulnerable species[6]. Additionally, as we show in Fig. 3, we can characterize geographically distinct populations of the same species. Past work has generally standardized each species’ flash patterns (at a particular temperature, usually 20°C [8], and across locations [7]), resulting in single, discrete measurements for the relevant statistics that are used as references for species identification in tandem with physical observation of captured specimens. Instead of relying on this, our work sets the new standard for representing each species’ subpopulations as their own entities with associated characteristics and variability. Future work can easily examine local climate effects in these different locations and further pair flash behavior to environmental effects.

Our automated trajectory detection method reliably measures the flash lengths and inter-flash gaps (Fig 3, Table 2) of firefly signals, especially as they are extracted from thousands of precise observations of the same species across multiple nights of filming. Human observers tend to over-estimate flash length (Fig 3C), which could be attributed to the reaction latency in a human observer between seeing the end of a flash and pressing a stopwatch. This problem is eliminated by our camera capture procedure, which is accurate to a frame resolution of 30 Hz, a significant improvement over measured human reaction times of order 0.2 s [25]. However, some limitations with the automated method exist. For example, it is possible that a given trajectory might not capture the entire flash pattern of an individual, due to overlapping signals from nearby fireflies, prolonged dark periods between series of successive flashes, and possible visual obstruction of blinks from the environment. Hence, it can be difficult to estimate the total count of flashes in a firefly’s signal, and the dark period between flash phrases is fully lost in most trajectories. This is mitigated in the characterization process by filtering only the classifier’s most probable flash patterns into a subset of data that should represent the best trajectories in the overall dataset for any given species. Combining our method with continued observations by human experts for the most complete characterization of any given species’ flashing behavior may be the optimal approach in the short term, especially in species known to have dark periods that exceed one second in length between flash phrases, or species that produce a single flash. However, the advancements presented here will indisputably accelerate this process and should only improve in effectiveness over time.

The model may additionally serve to answer future questions about the evolution of flash sequence distinguishability, especially under competing pressure to avoiding predation [26]. For example, high values off the diagonal in Fig. 2 represent relatively indistinguishable species. Species with high values on the diagonal are the most distinguishable. Correlating these results with geographical information may provide insight into character displacement and possibly improve the model. If species that are indistinguishable are more likely to be geographically isolated, their differentiability can be improved in our model with the additional application of location information.

Finally, this area is ripe for future development concerning scalability of the project and its applications in conservation biology. If, as the dataset gains more species, it becomes *more* difficult to discern between similar species, it is easy to pivot to an CRNN (convolutional recurrent neural network) structure [27], using spatial information extracted directly from the video recording to add additional features to the model that may serve to increase differentiability. As the capabilities increase, the classifier can be integrated into community-facing applications like iNaturalist [28], which in turn will enable faster data-training-classification loops and a widespread expansion of the model’s reach. Integration within community science is a vital component of the next steps: as discussed in [29], community monitoring programs have the potential to raise the public’s awareness and understanding about firefly endangerment and promote successful integration of policy and practice moving forwards. Our standardized, user-friendly data acquisition procedure can be adopted by volunteers and researchers to grow the dataset, with an eye toward recording flash sequences from data-deficient species.

## 4 Methods

### 4.1 Acquisition of flash sequence data

To extract flash sequence data, we perform 3D reconstruction of firefly swarms based on stereoscopic video recordings. Recordings were conducted at specific locations across the country where certain species were known to emerge [24]. Field researchers placed two spherical (360°) GoPro Max cameras at known distances from each other on a level surface (Fig. 1A). Recordings started at dusk when the first flashes were seen, and filmographers performed a simultaneous catch-and-release identification process to acquire ground-truth labels from visual inspection of the individuals present in the swarm. The movies were subsequently processed as described in a previous work [22, 24] to extract the 3D locations of flash occurrences. From these locations, we apply a simple distance-based linkage method to concatenate flashes into streaks and streaks into trajectories. We consider flashes at consecutive timesteps within a small radius to be part of the same streak; streaks occurring within both 1s and 1m of each other are assumed to come from the same individual and placed in a set of transitively connected streaks called a trajectory. To eliminate noise effects from the trajectory extraction, we threshold the trajectories to eliminate those that only contain one flash. The dataset [24] includes ten total species before the application of the thresholding process. Following the thresholding, we also remove any species from the dataset that have fewer than one hundred total trajectories, leaving us with seven species total. Finally, from the trajectories, we extract a binary time sequence by considering the time coordinates of flashes. This process enables the capture of individual behavior from simple footage of firefly swarms of any species, provided individuals of that species flash frequently enough to meet the threshold standards of the trajectory generation. Our data acquisition procedure highlights the presence of intraspecies behavioral variability, and characterizes this variability by representing flash patterns as distributions (Fig. 3A-C).

The result of this process is 58,916 flash trajectories from the seven species before thresholding, and 38,081 trajectories after those with only one flash have been removed. More than half of these are *P. carolinus* sequences - the majority class. About 1 percent of these are *P. forresti* and *P. bethaniensis* sequences - the minority classes. The rest of the classes are fairly evenly distributed. The dataset comprises binary sequences of between 14 and 1365 frames (0.467s to 45.5s) in length, each labeled with the corresponding firefly class.

### 4.2 Modeling

#### 4.2.1 Neural network architecture

RNNs are a class of neural networks suitable for sequence learning tasks. They are characterized by feedback connectivity and the consequent ability to encode long-range temporal dependencies, such as those intrinsic to firefly flash patterns. The defining computational step of an RNN is the hidden state update, which is a function of the input at the current timestep, *x*^(*t*)^, and the hidden state at the previous timestep, *h*^(*t–*1)^:

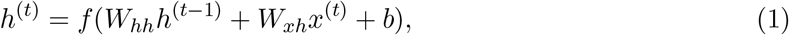

where *f* is a non-linear activation function, such as hyperbolic tangent, *W*_*hh*_ and *W*_*xh*_ are weight matrices that map hidden-to-hidden (i.e., the feedback connectivity) and input-to-hidden, respectively, and *b* is a bias term.

Importantly, Eq. 1 enables a recurrent neural network to ingest variable length input sequences, as the update rule can be applied recurrently, which is suitable for firefly flash sequences that are of variable temporal duration. However, if the input duration is sufficiently large–as is the case with some of flash pattern sequences–vanishing gradients will arise when computing weight updates via backpropagation through time (BPTT) [30] due to the non-linearity in *f*, ultimately prohibiting learning.

To address this issue, we leverage an extension to RNNs–gated recurrent units (GRUs)–that introduces gating mechanisms that regulate which information is stored to and retrieved from the hidden state [31]. These gating mechanisms enable the model to more effectively regulate the temporal context encoded in the hidden state and enables the encoding of longer-range temporal dependencies. Additionally, GRU RNNs are computationally more efficient than other kinds of RNNs like long short-term memory networks (LSTMs), and use fewer parameters, which was a consideration due to our plans for eventual downstream application of the model in real-time population monitoring. Consequently, we implement the model in PyTorch [32] as a 2-layer GRU with 128-dimension hidden layers, no dropout, and LeakyReLU activation layers with a negative slope of 0.1.

#### 4.2.2 Data preprocessing

To evaluate our model’s predictive ability on unseen data, we perform 60-fold stratified cross validation to ensure that each sequence in the dataset is used at least once in training and at least once in testing, but never simultaneously. Each fold divides the data into ninety percent training, and ten percent testing, preserving the same class ratios as the original dataset. Due to the severe class imbalance (e.g., some species only comprise 1% of the dataset, whereas others comprise close to 50% of the dataset), we perform random undersampling on the training set of each fold to equalize the class count in the training and validation sets for each fold. This takes the form of a secondary k-fold cross validation procedure to sample from each class until classes are equalized. All the remaining data are used for testing. The reported results are thus the ensemble precision, recall, and accuracy of each model on its respective test set of approximately 30,000 sequences, averaged over the 60 model folds. The ground truth targets are species names; we performed label encoding to transform the targets into machine-readable integers representing each class.

#### 4.2.3 Training and evaluation

We trained the model with the Adam optimizer [33] and evaluated performance via cross-entropy loss. During training, we set an early stopping callback that monitored the validation loss with a patience of 50 epochs to prevent overfitting on the training set. Additionally, to alleviate exploding gradients, we applied a gradient clipping of 0.1 to the gradient norms following the procedure recommended in [34]. We conducted a hyperparameter sweep over the batch size and learning rate, testing all combinations of batch size *∈* {8, 16, 32} and learning rate *∈* {10^−3^, 10^−4^, 10^−5^}. We selected the combination that had the highest validation set accuracy on a four-species subset of the data, which resulted in the choice of a batch size of 8 and a learning rate of 10^−5^. No data augmentation was applied during training.

We evaluate the performance of the RNN, along with the signal processing methods described in the following section, on the test data by examining the receiver-operating characteristic (ROC) curves for each species (Fig. 5A-E). Per-species precision and recall are tabulated in Fig. 5F.

### 4.3 Signal processing methods

For the purposes of comparison with the RNN, we implemented four alternative classifiers which use standard signal processing algorithms to compare our dataset against ground truth references for each species. We implement these classifiers using two types of ground truth references: “literature references”, which use flash patterns as previously published in the literature, and “population references”, which are generated by aggregating sequences in our own dataset.

#### 4.3.1 Literature references

“Characteristic” flash patterns for six out of the seven species analyzed in this paper, excluding *B. wickershamorum*, have been previously recorded and published in the literature. These recorded flash patterns hence served as the primary reference for researchers in identifying signals observed in the field. These reference flash patterns are typically reported pictorially; thus, we convert images to binary-valued time series by computing the relative sizes of flashes and gaps, in pixels. We determine the pixel-to-second conversion to then convert the sequence to a 30 frames per secod time series, matching the sampling frequency of our data.

These six reference time series then form the ground-truth comparisons against which our dataset is compared, using the four signal processing methods described below in section 4.3.3. We omit *B. wickershamorum* as there is currently no published reference pattern.

#### 4.3.2 Population references

We also generate “population references” by aggregating sequences in our own dataset. For each species, we first perform an 80:20 train:test split, similar to the preprocessing procedure performed for the RNN (see above section 4.2.2). The population references are obtained by averaging the sequences in each training set. The remaining test data is then classified using the signal processing algorithms described below in section 4.3.3. As with the RNN, we perform *N* = 100 iterations and take the ensemble average of the performance across all iterations.

#### 4.3.3 Signal processing algorithms

##### Jaccard index

The Jaccard index compares the similarity between two sets by taking the ratio of the size of the intersection of the sets with the size of the union [35] and has found broad application, for example in genomics [36]. For two binary-valued sequences 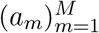 and 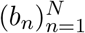of lengths *M* and *N*, respectively, with *a*_*m*_, *b*_*n*_ *∈* {0, 1} for all *m* and *n*, we define the size of the intersection as 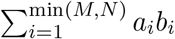, the number of ‘on’ (flashing) bits that occur simultaneously in both sequences. We define the union as 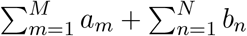, the number of on bits for both sequences combined. The Jaccard index can also be evaluated in the same manner for two sequences that are binary-valued. Generally speaking, the intersection can also be thought of as the dot product between the two sequences. To classify a sequence using the Jaccard index, the Jaccard index between a sequence and each species reference is computed, and the softmax of the vector of Jaccard index values is computed to determine a probability of the sequence being from each species. The predicted species is then the argument maximum (arg max) of the softmax vector.

##### Dot product

The dot product between two sequences is given by the sum of the product of the sequences, i.e., 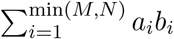 for two sequences 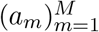 and 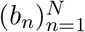 of lengths *M* and *N*, respectively. Sequences are then classified by taking the arg max of the softmax of the dot product with each reference.

##### Dynamic time warping

Dynamic time warping (DTW) is an algorithm that computes the distance between two timeseries by locally stretching or compressing the sequences to optimize the match. DTW is useful for comparing timeseries that are qualitatively similar but vary locally in speed and has found significant application in speech recognition [37–40]. We implement DTW in MATLAB [41] to compute the distance between sequences and species references. Similarly to the other metrics, the predicted species is taken to be the arg max of the softmax of the distances, which is a vector of probabilities that maps to the probability of each label.

##### SVM

Each flash sequence can be parametrized by 3 values (Fig. 1A): the number of flashes in the sequence, the average duration of each flash, and the average length of spaces between flashes (inter-flash gap). We perform support vector machine (SVM) classification in this 3-dimensional space, using a radial basis kernel function.

### 4.4 Characterization

The data acquisition procedure is not without noise, so we perform filtering to produce accurate quantitative characterization of flash phrases that falls in alignment with previous literature observations. We leveraged the ability of the RNN to distinguish between sequences by choosing the sequences which the RNN scored highest as the top one hundred most confident classifications for each species. This subset acts as the dataset on which characterization exercises are performed for Fig. 1B and Fig. 3. The procedure is as follows:

1. Initialize the empty list *c*
2. for each *i* sequence in the test set *D* :
  (a) Run a forward model step
  (b) Let *p* = the maximum probability in the resulting vector of softmax predictions
  (c) If the index of *p* corresponds with the correct label, add the pair (*p*, index) to the list *c*
3. Sort *c* by probability *p* and choose the top 100
4. Index into the dataset D using the associated indices of the top 100 probabilities to produce the subset Characterizing in this way leverages the variability in the entire dataset by training the predictive classifier, then asks the predictive classifier only for what it is most confident about in order to filter out sequences that may be missing flashes or exhibiting patterns that are far from the statistical norms of the species.

## 5 Acknowledgements

The authors would like to thank many collaborators on this project. Avalon Owens, Paul Shaw, Richard Joyce, Cheryl Mollahan, Peggy Butler, Jason Davis, and Julie Hayes all were excellent volunteer data gatherers following our methodology. Candace Fallon and Sara Lewis were invaluable allies in consultation about firefly behavioral ecology and conservation threats. We’d also like to thank Airy Gonzalez Peralta, Yuti Gao, and Michael Hoffert for their insight and feedback as the project developed, and Brett Melbourne for his patience and helpful guidance through endless papers related to machine learning in ecology as we moved towards the right classification approach. Finally, the authors would like to thank Golnar Gharooni Fard, Nolan Bonnie, and Sam Pugh for thoughtful discussion and insights about the paper. O.P. acknowledges support from the National Geographic Society grant NGS-84850T-21, the Research Cooperation for Science Advancement grant 28219, and the BioFrontiers Institute at the University of Colorado Boulder.

## 5.1 Author contributions

O.P. conceived the study and provided funds to conduct research. O.P., C.N., O.M., M.L.I., and D.M.N. designed the research. R.S. and O.P. provided flash sequence data. M.L.I. and D.M.N. prototyped preliminary models. O.M. and C.N. analyzed the data, built the final models, and performed statistical analyses. M.C. and R.L. provided signal processing and machine learning algorithmic insights. O.M., C.N., R.S., M.C., R.L., M.L.I., and O.P. wrote the paper. O.P. and R.L. supervised the project.

## 5.3 Data and materials availability

The associated code and data files for this project can be found here: https://github.com/peleglab/FireflyClassification. Following instructions in the associated README.md should allow easy regeneration of the figures and replication of the workflows described in this paper. All data needed to evaluate the conclusions in the paper are present in the github repository, the Supplementary Information, and/or the associated dataset paper [24].

## Supplementary Information

### 5.4 Batch effects

The dataset was assembled by several scientists operating at different locations across different years, so it is important to consider the possibility that the network is isolating a batch-related feature. We control for this in several ways: first, the methodology for video collection is standardized following procedural guidelines outlined in [24]. These are easy to follow, clearly delineated, and often quality checked before the data are gathered. Second, the videos are post-processed to transform them into binary sequences, eliminating extraneous factors that may normally contribute to batch effects in neural network training. Third, the resulting sequences are all transformed to the same time scale to avoid preference given to a certain frame rate. We believe these steps are sufficient to remove any noticeable effect generated by minor differences in data collection moving forward, but testing this is important to make sure the model is not learning a characteristic of the sequences that is isolated to a certain data gatherering event. To test this, we performed 5-fold cross-validation on the recurrent neural network where all of the data for a particular data gatherer was held out of the training set. For each fold, we isolated all of the *P. frontalis* and *P. carolinus* sequences in the dataset coming from R.S. [24]. We did this because *P. carolinus* and *P. frontalis* are two of the species for which there are multiple data-gathering events and camera operators present in the dataset, and R.S. took part in some of the data collection for each of these species. Holding their data out from the training dataset means the model will not be able to see any of those sequences, so if there is a batch effect, it should not be able to recognize them. Then for each of the 5 folds, we selected a different fold of the remaining *P. carolinus* and *P. frontalis* data to be part of the training set. The other species were treated normally, and the results are in Fig. S1. It is clear that there is no batch effect, as the model performs just as well during the 5-fold cross validation as it does without any data holdout. This is likely thanks to the distinctive flash length of *P. frontalis*; additionally, despite the differences *between* distributions of the inter-flash gap for *P. frontalis*, the overall distribution is still uniquely situated between 0.5s-0.8s, which is unlike any of the other species in the dataset. This analysis supports the ability for our model to scale with different volunteers acting in different locations serving as the data-gatherers, and eventually to automated camera setups, without concern.

**Figure S1:**
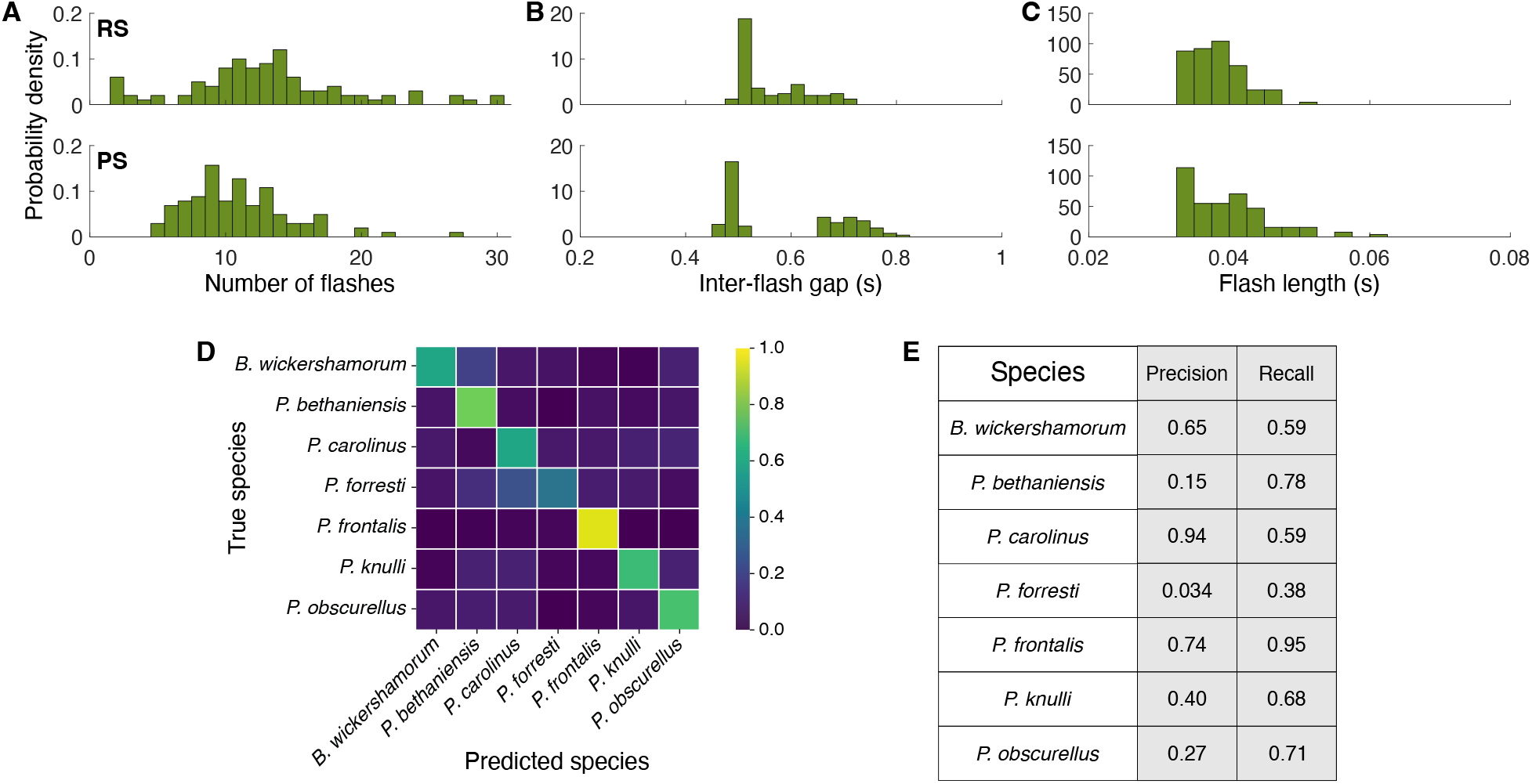
Quantification of batch effects. **A-C:** Probability distributions of flash parameters from *P. frontalis* sequences collected by two different data gatherers, labeled RS (top) and PS (bottom). The distributions for number of flashes **(A)** and flash length **(C)** are similar, while the inter-flash gap **(B)** distributions are notably different due to different temperatures during data collection for the two different gatherers. **D:** Confusion matrix for 5-fold cross validation of RNN where all *P. frontalis* and *P. carolinus* data gathered by R.S. is held out from training. The RNN achieves similar performance to the model described in the main text, which was trained on both RS- and PS-collected *P. frontalis* data. **E:** Table of per-species precision and recall during batch effect cross-validation.

### 5.5 Literature ROC results

The literature methods perform poorly compared to population reference methods, with the SVM achieving the best result (precision = 0.32, recall = 0.28). However, we observe that the SVM classifier predicts most samples as either *P. bethaniensis* or *P. forresti*, The receiver-operating characteristic (ROC) curves for the literature-based methods are shown in Fig. SI2, along with the per-species precision and recall. All four methods perform poorly on *P. carolinus*. SVM and DTW result in near-perfect recall on *P. knulli*, but poor precision; conversely, these methods give near-perfect precision on *P. frontalis*, but poor recall.

These results reveal that the existing literature-based reference patterns fall short in accurately representing our collected dataset. This may be due to the different conditions under which the literature-based references; moreover, by creating population references, we can better capture the behavioral variability found in the data.

**Figure S2:**
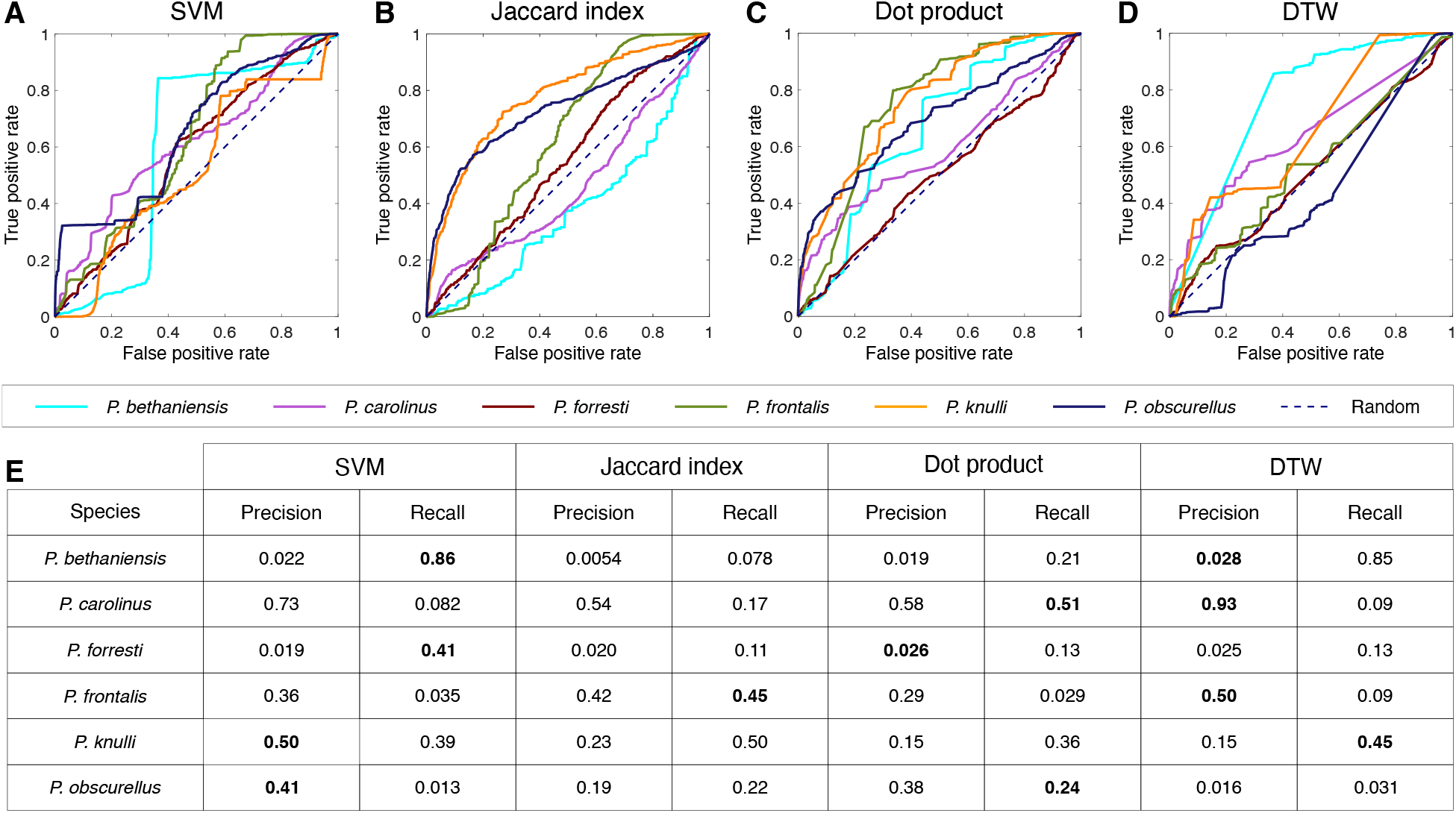
Classification results for literature-based references. A-D: Receiver operating characteristic (ROC) curves representing the true positive rate (TPR) as a function of the false positive rate (FPR) across all model thresholds of classification, labeled by method. E: Table of per-species precision and recall.

